# Custom long non-coding RNA capture enhances detection sensitivity in different human sample types

**DOI:** 10.1101/2021.04.14.439879

**Authors:** Annelien Morlion, Celine Everaert, Justine Nuytens, Eva Hulstaert, Jo Vandesompele, Pieter Mestdagh

## Abstract

Long non-coding RNAs (lncRNAs) are a heterogeneous group of transcripts that lack protein coding potential and display regulatory functions in various cellular processes. As a result of their cell- and cancer-specific expression patterns, lncRNAs have emerged as potential diagnostic and therapeutic targets. The accurate characterization of lncRNAs in bulk transcriptome data remains challenging due to their low abundance compared to protein coding genes. To tackle this issue, we describe a unique short-read custom lncRNA capture sequencing approach that relies on a comprehensive set of 565,878 capture probes for 49,372 human lncRNA genes. This custom lncRNA capture approach was evaluated on various sample types ranging from artificial high-quality RNA mixtures to more challenging formalin-fixed paraffin-embedded tissue and biofluid material. The custom enrichment approach allows the detection of a more diverse repertoire of lncRNAs, with better reproducibility and higher coverage compared to classic total RNA-sequencing.

## Introduction

While the majority of the human genome is actively transcribed into RNA transcripts, most of these transcripts do not code for proteins (Djebali *et al.*, 2012). The non-coding RNA transcripts longer than 200 nucleotides belong to the heterogeneous group of long non-coding RNAs (lncRNAs), half of which are not poly-adenylated (Lorenzi *et al.*, 2019). These lncRNAs are known to influence gene expression at both the transcriptional and post-transcriptional level through a variety of mechanisms (Mercer *et al.*, 2009; Robinson *et al.*, 2020). Moreover, lncRNAs often show a particular cell- or cancer-type specific expression pattern (Iyer *et al.*, 2015), which adds to their biomarker potential.

In the past, several high-throughput methods have been developed to profile the long non-coding RNA transcriptome, study their structure or define their function (Cao *et al.*, 2019; Turner *et al.*, 2019). Because of their generally low abundance compared to protein coding genes, quantification of lncRNAs in bulk transcriptome data remains challenging. Enrichment strategies favoring lncRNAs over the more abundant mRNAs could therefore result in more lncRNAs being detected with a better transcript coverage, improving downstream analysis. A promising method is RNA capture sequencing, a short-read sequencing method that can enrich RNA targets of interest using oligonucleotide probes that are specifically designed to tile the target sequences. These RNA capture sequencing technologies have mainly been applied for deep sequencing of a selection of lncRNAs (Mercer *et al.*, 2014; Clark *et al.*, 2015). Recently, the GENCODE consortium extended this method by applying long-read sequencing after capturing about 14,470 lncRNAs genes to improve their structural annotation (RNA Capture Long Seq, RNA CLS) (Lagarde *et al.*, 2017).

In this study, we describe a custom lncRNA capture sequencing approach that targets a very comprehensive human lncRNome. This custom capture approach was evaluated on various sample types ranging from high-quality RNA mixtures to more challenging formalin-fixed paraffin-embedded (FFPE) tissue and biofluid material.

## Material and methods

### Probe design

Probes were designed against the highly confident set of LNCipedia 5.2 (hg19 genome build). First, extended exons were created by concatenating each set of overlapping exons. For each of these extended exons, probes of 120 nucleotides were tiled, resulting in (number of nucleotides)-119 probes per concatenated exon. These exon tiling probes were mapped against repeat regions and protein coding genes to filter out these that would capture off-target fragments.

The resulting probe pool was extended with probes designed to capture both the Sequin and ERCC spikes. These probes are 120-mers designed by tiling the spike-in sequences and, inherent to the spike-design, these do not align to the human genome.

Further filtering was done by retaining the 120-mers with a GC content between 25-70%, a GC-based Tm between 60-80 °C and a ΔG larger than -7 (calculated by UNAFold (version 3.8) settings: hybrid-ss-min -E -n DNA -t 54 -T 54). The remaining probes underwent a selection aimed at obtaining the minimal number of probes for an optimal coverage. In total, 565,878 probes against LNCipedia, 81,089 probes against novel genes (not discussed in this paper) and 2427 spike-in RNA probes were retained. Probes were synthesized by Twist Biosciences.

### Sample collection and RNA purification

Sample collection was approved by the ethics committee of Ghent University Hospital, Ghent, Belgium (#B670201734450 and #B670201733701) and written informed consent was obtained from all donors. FFPE tissues were obtained from two colon cancer patients; the biofluid samples (seminal and blood plasma) were collected from healthy donors.

### Platelet depleted blood plasma

Venous blood from two healthy donors was drawn from an elbow vein after disinfection with 2% chlorhexidine in 70% alcohol. All blood draws were performed with a butterfly needle of 21 gauge (BD Vacutainer, Push Button Blood Collection Set, #367326, Becton Dickinson and Company, NJ, USA) and blood was collected in 10 ml BD Vacutainer K2-EDTA tubes (#367525, Becton Dickinson and Company, NJ, USA). The tubes were inverted 5 times and centrifuged immediately after blood draw (15 min at 2500 g, room temperature, without brake). Per donor, the upper plasma fractions were pipetted (leaving approximately 0.5 cm plasma above the buffy coat) and pooled in a 15 ml tube. After gently inverting, the pooled plasma fraction was centrifuged again (15 min at 2500 g, room temperature, without brake) and the upper fraction was transferred to a new 15 ml tube, leaving approximately 0.5 cm plasma above the separation. The resulting platelet depleted plasma was gently inverted, snap-frozen in five aliquots (Safe-Lock cup DNA LoBind 2 ml PCR clean tubes, Eppendorf, #0030108078) and stored at -80 °C. Platelets were counted and the degree of hemolysis was determined by measuring levels of free hemoglobin by spectral analysis using a NanoDrop 1000 Spectrophotometer (Thermo Fisher Scientific). The entire plasma preparation protocol was finished in two and a half hours. 200 μl was used for each RNA isolation.

### Seminal plasma

Semen samples of healthy donors were produced by masturbation into a sterile container and were allowed to liquefy for 30 min at 37 °C. Samples were centrifuged to remove contaminating cells (10 min at 2000 g, room temperature, without brake) and stored at -80 °C within two hours after collection. 200 μl was used for each RNA isolation.

### Biofluid RNA purification

RNA was isolated with the miRNeasy Serum/Plasma Kit (Qiagen, #217184) according to the manufacturer’s instructions. An input volume of 200 μL was used for all samples. Per 200 μL biofluid input volume, 2 μL sequin spike-in controls (Garvan Institute of Medical Research) were added before RNA isolation, in a 1/1300 000 dilution to blood plasma and in 1/1300 dilution to seminal plasma. Total RNA was eluted in 12 μL of RNAse-free water for (blood) platelet depleted plasma, and in 20 μL of RNAse-free water for seminal plasma – in order to adjust for viscosity. After RNA isolation, 2 μl External RNA Control Consortium (ERCC) spikein controls (ThermoFisher Scientific, #4456740) were added to the RNA isolation eluate of blood plasma and seminal plasma in a dilution of 1/1000 000 and 1/1000, respectively. gDNA heat-and-run removal was performed by adding 1 μl HL-dsDNase (ArcticZymes #70800-202, 2 U/μl) and 1.4 μl reaction buffer (ArcticZymes #66001) to the combination of 12 μl RNA eluate and 2 μl ERCC spikes, followed by an incubation of 10 min at 37 °C and 5 min at 58 °C. RNA was stored at -80 °C and only thawed on ice immediately before the start of the library prep. Multiple freeze-thaw cycles did not occur. RNA obtained from three RNA isolations was pooled per biofluid and per sample to avoid RNA isolation induced variation. This pooled RNA was used as starting material for the different library preparations.

### FFPE

Tumor RNA was isolated from five 10 μM sections of a formalin-fixed paraffin embedded (FFPE) tissue block, applying macrodissection based on histopathological evaluation of hematoxylin and eosin stained slides to select regions with high tumor cellularity. Within two days after sectioning, the tissue sections were scraped into microcentrifuge tubes, centrifuged for 5 min at 20,000 g, and deparaffinized in 320 μl Deparaffinization Solution (Qiagen, #19093) for 3 min at 56 °C on a thermomixer (500 rpm). Samples were then cooled to room temperature for 15 min. Subsequently, RNA was isolated using the miRNeasy FFPE Kit (Qiagen, #217504), according to the manufacturer’s protocol. gDNA heat-and-run removal was performed by adding 1 μl HL-dsDNase (ArcticZymes #70800-202, 2 U/μl) and 0.68 μl reaction buffer (ArcticZymes #66001) to 6.82 μl RNA (100 ng), followed by an incubation of 10 min at 37 °C and 5 min at 58 °C.

### MAQCA/B

Two commercially available RNA samples, MAQCA and MAQCB, were used. MAQCA is the Quantitative PCR Human Reference Total RNA (#750500, Agilent technologies), extracted from cell lines representing different human tissues. MAQCB is FirstChoice Human Brain Reference RNA (#AM7962, Life Technologies). gDNA heat-and-run removal was performed on both RNA samples by adding 1 μl HL-dsDNase (ArcticZymes #70800-202, 2 U/μl) and 0.68 μl reaction buffer (ArcticZymes #66001) to 6.82 μl RNA (100 ng), followed by an incubation of 10 min at 37 °C and 5 min at 58 °C.

### Library preparation

After RNA purification, four libraries were prepared for each sample: two technical replicates for total RNA-seq and two technical replicates for custom lncRNA capture sequencing.

### SMARTer Stranded Total RNA library preparation

Sequencing libraries were generated using SMARTer Stranded Total RNA-Seq Kit v2 - Pico Input Mammalian (Takara Bio, #634413). The library preparation protocol started from 6 μL eluate for the biofluid samples and 100 ng (or 10 ng) RNA for FFPE and MAQC. The recommended amount of input RNA for SMARTer Stranded Total RNA sequencing is only up to 10 ng while the capture method uses 100 ng. To make sure our analyses were not biased, we decided to use the total RNA-seq method with 100 ng RNA input as well but also included 10 ng input samples. As shown in SFig 6 the results of 10 vs 100 ng RNA are similar. LncRNAs that are only detected using one of the input amounts are mostly low abundant lncRNAs that are just below the threshold. Compared to the manufacturer’s protocol, the fragmentation step was set to 2 min at 94 °C, hereafter the option to start from high-quality or partially degraded RNA was used. During the final RNA seq library amplification, 16 PCR cycles were used for the samples derived from platelet depleted (blood) plasma, 12 PCR cycles were used for the other samples, and the cycles were followed by an extra 2 min at 68 °C before cooling them down to 4 °C. Library quality control was performed with the Fragment Analyzer high sense small fragment kit (Agilent Technologies, sizing range 50 bp-1000 bp). As Fragment Analyzer profiles showed the presence of multiple adapter dimers, the final AMPure Bead Purification step was repeated (17 μl AMPure beads added to each sample - 20 μl Tris Buffer was used to resuspend the beads – and elution volume of 18 μl).

### Custom RNA capture library preparation

Custom RNA capture-based libraries were prepared starting from 8.5 μL eluate for biofluid samples and 100 ng RNA for FFPE and MAQCA/B using the TruSeq RNA Exome Library Prep Kit (Illumina, USA). Library preparation happened according to the manufacturer’s protocol with some minor modifications. Fragmentation of RNA with the thermal cycler was set for 2 min at 94 °C (instead of 8) and incubation to synthesize first strand cDNA for 30 min at 16 °C (instead of 60 min). After library validation with Fragment Analyzer (Agilent Technologies), the Twist Human Core Exome EF Multiplex protocol (Twist Bioscience, San Francisco, USA) was used starting with the pooling of amplified indexed libraries in sets of eight. One pool consisted of MAQCA/B and seminal plasma libraries (with the required 187.5 ng per sample), the other pool was a low-input pool containing the FFPE and (blood) plasma libraries (with the available 20 ng per sample). Heated hybridization mix was added to the custom capture probes without cooling down to room temperature in order to prevent the probes from precipitating. After hybridization of probes with pools and binding to streptavidin beads, post capture PCR amplification was performed at 8 cycles for the high-input pool and 12 cycles for the low-input pool. After cleanup, the final libraries were validated with Fragment Analyzer (Agilent Technologies).

### Sequencing

Based on qPCR quantification with the KAPA Library Quantification Kit (Roche Diagnostics, #KK4854), samples were pooled and loaded on NextSeq 500 with a loading concentration of 1.6 pM for the custom RNA capture libraries and 1.3 pM for the SMARTer Stranded Total RNA libraries. Paired end sequencing was performed (2 x 75 nucleotides). Custom RNA capture sequencing resulted in 168 million PE reads (median: 8.4 million PE reads/sample), SMARTer Stranded Total RNA sequencing resulted in 110 million PE reads (median: 10.5 million PE reads/sample). FASTQ data is currently being deposited in EGA.

### Sequencing data quality control

The SMARTer Stranded Total RNA seq libraries were trimmed using cutadapt (v.1.16) to remove 3 nucleotides of the 5’ end of read 2 (Martin, 2011). Reads with a low a base calling accuracy (< 99% in at least 80% of the bases in both mates) were discarded. To enable a fair comparison, we started data-analysis from an equal number of reads by downsampling to the minimum available paired-end reads per sample type (rounded to half a million): 6.5 million for FFPE, 7.5 million for MAQCA/B, 6 million for seminal plasma, 3 million for platelet-depleted (blood) plasma. Downsampling was done with Seqtk (v1.3) (Li, 2021). Next, read duplicates were removed with Clumpify (BBMap v.38.26, standard settings) using the following specifications: paired-end mode, 2 substitutions allowed, kmersize of 31, and 20 passes (Bushnell, 2021). For duplicate removal, only the first 60 nucleotides of both reads were considered to account for the sequencing quality drop at the end of the reads. Full-length read sequences were retrieved after duplicate removal for further quantification.

### Quantification of Ensembl and LNCipedia genes

Strand-specific transcript-level quantification of the deduplicated FASTQ files was performed with Kallisto (v.0.44.0) in –rf-stranded mode (Bray *et al.*, 2016). Quantification was performed with two references. The first one is a custom Ensembl v75 reference where lncRNAs are only taken from LNCipedia 5.2 (high-confidence set) (Volders *et al.*, 2019; Yates *et al.*, 2020). This reference was used to design the custom probes. The second reference is only based on a more recent version of Ensembl v91.

Further processing was done with R (v.4.0.3) making use of tidyverse (v.1.3.0). A count threshold for filtering low abundant genes was set based on an analysis of single positive genes in technical replicates (Mestdagh *et al.*, 2014). Single positives are genes with a zero count value in one replicate and a non-zero value in the other one. After applying a threshold of 10 counts, at least 95% of the single positives are removed (SFig 3).

## Results

In brief, 565,878 lncRNA capture probes of 120 nucleotides in length were designed against the high confidence set of LNCipedia v5.2 (Volders *et al.*, 2019) that comprises 107,039 transcripts belonging to 49,372 lncRNA genes. This probe set targets 45,284 lncRNA genes or 91.72% of the LNCipedia high confidence set. The median number of probes designed per lncRNA is 5 (SFig 1a), ranging from 1 up to 1675 probes for lnc-TBC1D22A-4 (with a length of 152,544 bp). The selected probe designs have a median GC of 43.33%, a Tm of 72.42 °C and ΔG of -2.8 (SFig 1b,c,d).

The custom lncRNA capture approach was applied to RNA from four different human sample types: high-quality RNA (artificial RNA mixture from human cell lines, MAQCA, and human brain reference RNA, MAQCB (Shi *et al.*, 2006)), formalin-fixed paraffin embedded colon tissue samples (FFPE), platelet-depleted blood plasma and seminal plasma. Each sample was also profiled with a total RNA-sequencing workflow representing the gold standard for quantification of both polyadenylated and non-polyadenylated lncRNAs.

We observed a clear enrichment of the lncRNA fraction with custom lncRNA capture compared to total RNA-seq when mapping reads to a LNCipedia transcriptome reference. Up to 75% of mapped reads in the custom capture method are derived from lncRNAs (Fig 1a), which is a 3.5-fold enrichment compared to total RNA-seq for FFPE, 4-fold for high quality MAQCA/B RNA and 8.5-fold for seminal plasma. This enrichment was also observed when aligning reads to a less comprehensive lncRNA reference (the Ensembl v91 reference), although the level of enrichment was lower (SFig 2a). In blood plasma, only a small fraction of reads aligned to lncRNAs for both the custom lncRNA capture and total RNA-seq method, resulting in the detection of just a few hundred lncRNAs (data not shown). In FFPE, the fraction of reads mapping to ribosomal RNA was higher in total RNA-seq (38% and 52% for donor 1 and 2, respectively) compared to custom capture (10% and 20% for donor 1 and 2, respectively) (Fig 1a). In other sample types, the lower fraction of lncRNA reads in total RNA sequencing compared to custom capture sequencing is almost exclusively compensated by a higher fraction of protein coding RNA (mRNA) reads.

**Fig 1:**
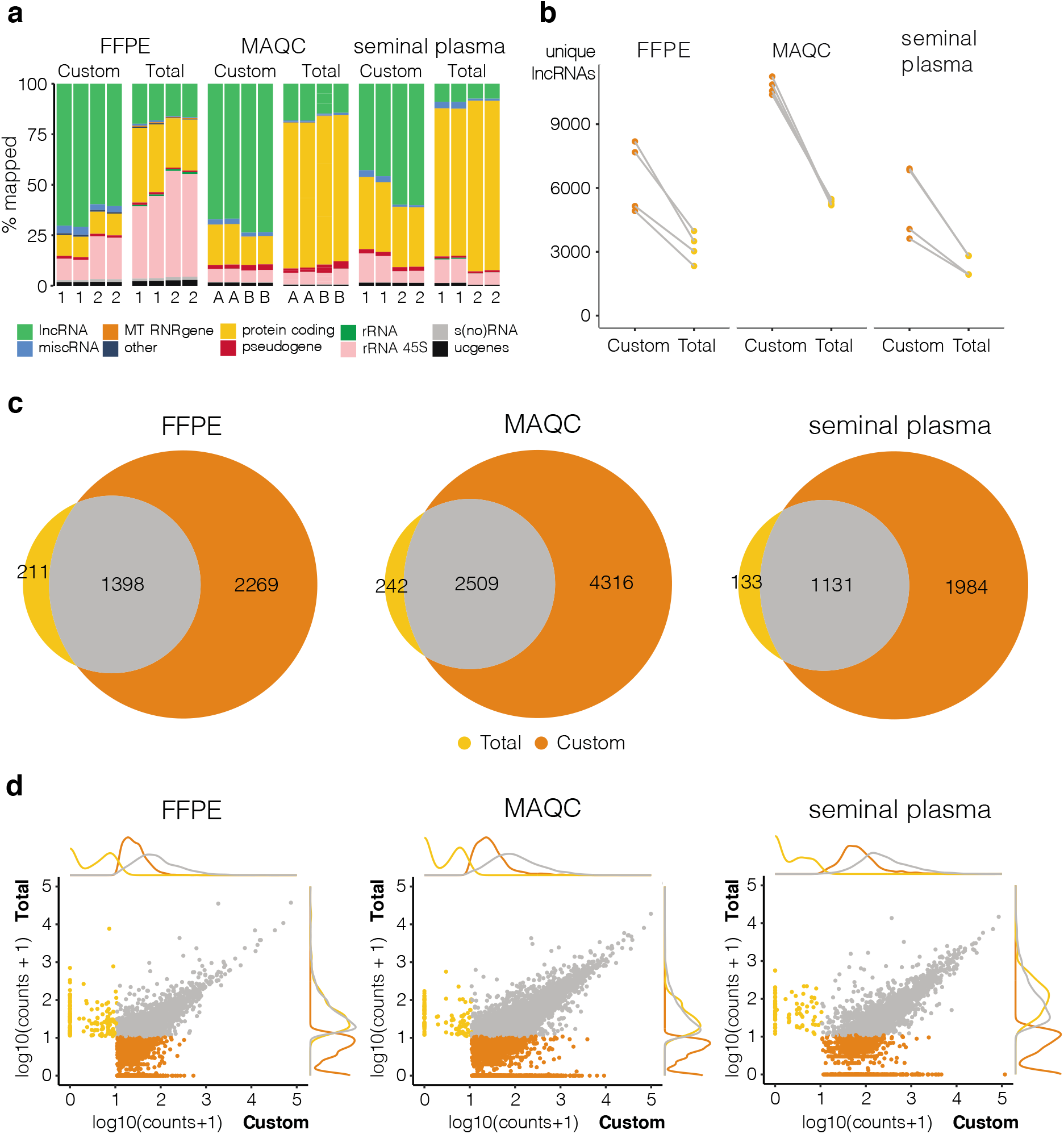
Custom capture sequencing (Custom) is able to detect more lncRNAs than total RNA-sequencing (Total). Quantification based on combined reference of Ensembl and LNCipedia. a: RNA biotype distribution plot of mapped reads where 1 and 2 indicate the two different donors and A and B refer to MAQCA and MAQCB, respectively (lncRNAs: high-confidence lncRNAs based on LNCipedia 5.2; miscRNA: miscellaneous RNA, non-coding RNA that cannot be classified; MT RNR gene: mitochondrially encoded ribosomal RNAs; protein coding: protein coding RNA transcripts; pseudogene; rRNA (45S): (45S) ribosomal RNA; s(no)RNA: small nuclear/nucleolar RNA; ucgenes: unannotated cancer genes; other: T cell receptor genes, Immunoglobulin genes, TEC (To be Experimentally Confirmed) - regions with EST clusters that have polyA features that could indicate the presence of protein coding genes, vaultRNA - short non coding RNA genes that form part of the vault ribonucleoprotein complex; microRNAs; ribozymes); b: number of unique lncRNAs with at least 10 counts (filter threshold), data points from same donor or MAQC type are linked (grey lines); c: overlap between lncRNAs that are detected above threshold in all replicates of a certain library prep method, plots made with eulerr package (v6.1.0) in R; d: correlation and density plots of overlapping (grey) and specific lncRNAs for custom capture (orange) and total RNA- sequencing (yellow); lncRNAs below count threshold in both methods were left out.

After downsampling to the same number of reads, we applied a minimal coverage of 10 counts to select for lncRNAs that are reproducibly detected (SFig 3) and compared detection sensitivity between both methods. Although both methods were able to detect several thousands of lncRNAs, the custom capture method on average resulted in two times more uniquely detected lncRNAs compared to total RNA-seq (Fig 1b). The maximum number of detected lncRNAs with the custom capture approach was 8186 for FFPE, 11,238 for MAQCA/B, and 6910 for seminal plasma. As expected, the majority of lncRNAs detected in all total RNA-seq replicates were also detected in all custom capture replicates: 87%-91% of lncRNAs based on LNCipedia reference (Fig 1c); 83%-91% based on Ensembl reference (SFig 2c). More importantly, custom capture enabled the detection of several thousands of additional lncRNAs (59%-61% of all lncRNAs reproducibly detected by custom capture were not detected by total RNA-seq), illustrating the sensitivity of this procedure (Fig 1c & SFig 2c). Expression abundance analysis revealed that these uniquely detected lncRNAs are generally less abundant compared to lncRNAs detected by both methods (Fig 1d & SFig 2d).

Next, we evaluated reproducibility based on absolute log2 fold changes of lncRNA abundance between technical replicates (ideally, these fold changes are close to zero). As shown in Fig 2 and SFig 4, we observed a higher fraction of lncRNAs with a log fold change close to zero in the custom capture approach compared to the total RNA-seq approach, indicating a better reproducibility for the custom capture approach. Only the total RNA-seq data of seminal plasma from donor 1 showed better reproducibility (SFig 4e), yet the custom approach in general still had lower fold changes between technical replicates (Kolmogorov-Smirnov test p-value < 0.001). Note that seminal plasma from donor 1 also resulted in a lower number of unique lncRNAs than that of donor 2 (Fig 1b).

**Fig 2:**
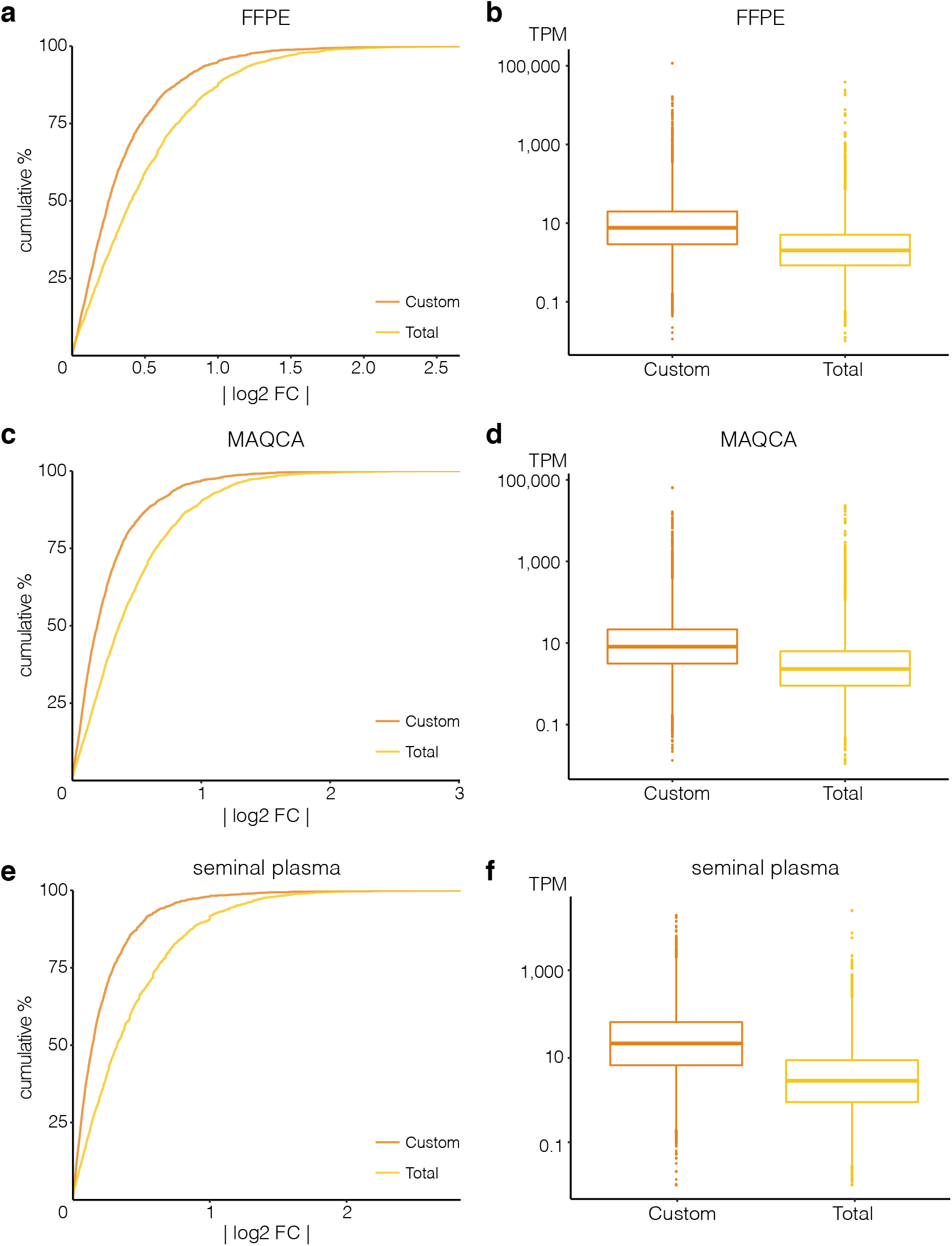
Custom capture seq (Custom) has a higher lncRNA count reproducibility and coverage than total RNA-seq (Total). Cumulative distributions of absolute log2 fold changes (log2 FC) between lncRNA counts in the two technical replicates are shown for (a) FFPE from donor 1, (c) MAQCA, and (e) seminal plasma from donor 2. Kolmogorov–Smirnov tests each time showed significant difference in distributions between Total and Custom (p-value < 0.001). Boxplot of corresponding transcripts per million (TPM) values of these lncRNAs are shown in (b) for FFPE, (d) for MAQCA, and (f) for seminal plasma.

We also compared transcript coverage of lncRNAs that were detected with both approaches by looking at their TPM distributions. In general, coverage was higher in the custom capture approach than in total RNA-seq (Fig 2 & SFig 4). Median TPM values for the custom capture and total RNA-seq approach, respectively, were 8.2 and 2.0 TPM in FFPE, 9.7 and 2.5 TPM in MAQCA/B, and 16.1 and 2.3 TPM in seminal plasma. In terms of gene body coverage, both methods covered the entirety of the lncRNA body with an expected lower coverage towards the 5’ and 3’ ends. The custom capture sequencing, however, showed a more pronounced reduction in coverage towards the 3’ end of the lncRNAs compared to total RNA-seq (SFig 5). Finally, we wanted to further assess the relevance of the custom capture approach for biological or clinical applications. We evaluated the abundance of previously described prostate-cancer related lncRNAs (Helsmoortel et al., 2018) in seminal plasma samples between both methods. As shown in Fig 3, coverage of detected lncRNAs is consistently higher with custom capture sequencing than with total RNA-seq. In total, 16 prostate-cancer related lncRNAs were detected above threshold in at least one sample. While none of those lncRNAs were exclusively detected by total RNA-seq, five lncRNAs (LINC01564, lnc-HNF1A-1, lnc-SPATA31A6-6, PCA3, and PCAT7) were detected by custom capture sequencing only. The custom capture counts of these lncRNAs ranged from 11 to 40 when taking the mean of both technical replicates of donor 2. Yet, these lncRNAs (except LINC01564) did not reach the detection threshold in custom capture sequencing samples of donor 1. For the 11 lncRNAs that were detected with both methods, custom capture sequencing resulted in 2 to 15 times more counts compared to total RNA-seq with an average fold change of 6. This increased sensitivity could greatly benefit biomarker research.

**Fig 3:**
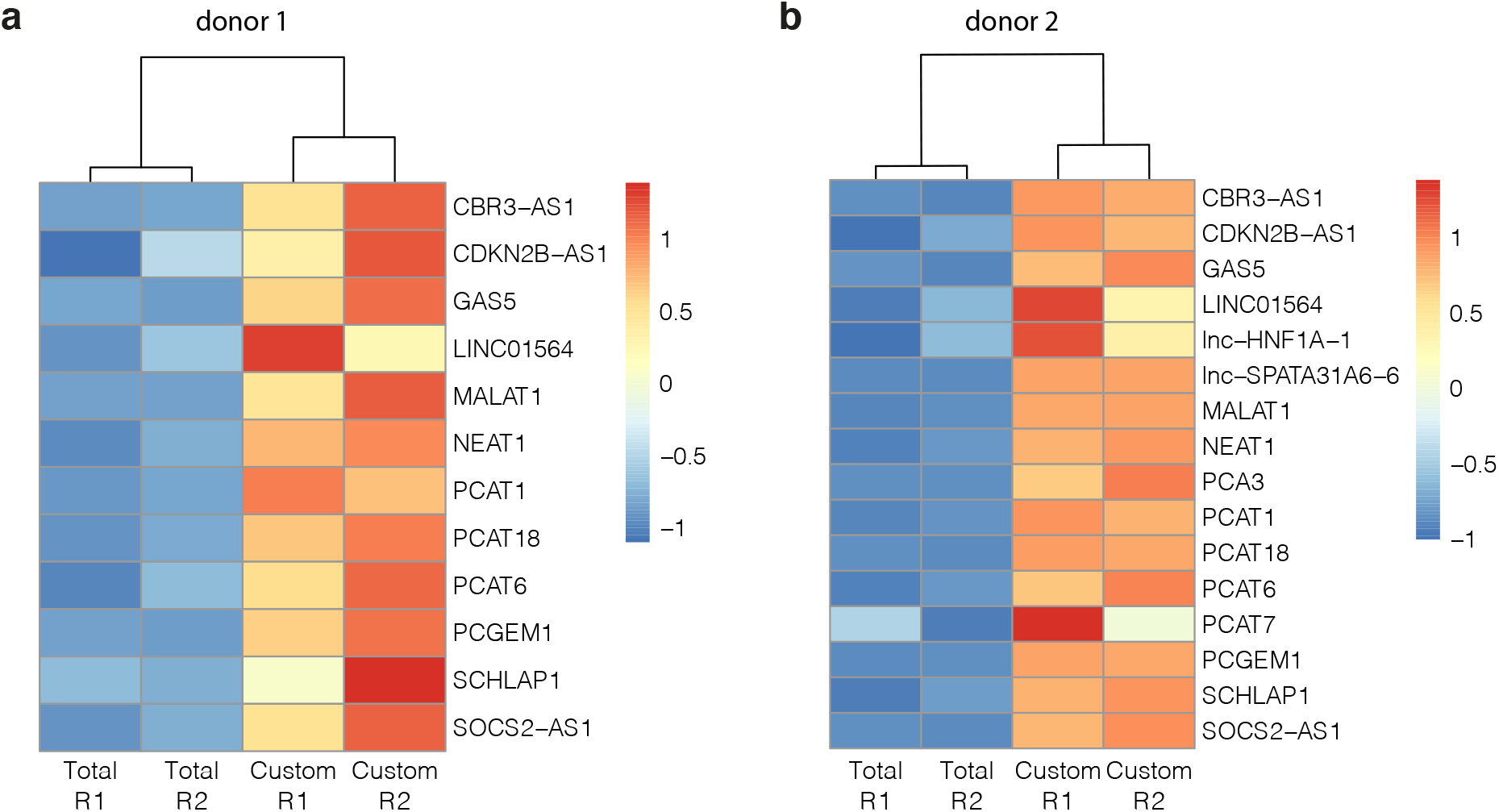
Higher coverage for prostate-cancer related lncRNAs with custom capture (Custom) than total RNA-sequencing (Total). Heatmaps based on z-score transformed lncRNA counts of seminal plasma samples from donor 1 (a) and donor 2 (b), respectively. Per donor, only lncRNAs detected above count threshold (10 counts) in at least one replicate were considered. A higher z-score (orange/red) indicates relatively more coverage. Complete clustering of samples based on Euclidean distance. R1: technical replicate 1; R2: technical replicate 2.

In summary, these findings demonstrate the added value of our custom lncRNA capture method for applications aimed at establishing a more complete lncRNA expression landscape.

## Discussion

An extensive enrichment combined with a higher coverage of lncRNAs may further improve our understanding of lncRNA association to various conditions or phenotypes. Studies aiming to identify lncRNA biomarkers could equally benefit from these advantages. We have demonstrated a superior performance of custom lncRNA capture sequencing compared to classic total RNA-sequencing, across different sample types. Fewer reads are consumed by RNA biotypes other than lncRNAs, which results in a better lncRNA coverage. Interestingly, we also observed a better lncRNA detection reproducibility between technical replicates for the custom capture compared to total RNA-seq (Fig 2). Deeper sequencing could event further improve the performance.

The custom capture method, however, did not outperform the total RNA-sequencing method in platelet-depleted blood plasma. In these samples, both methods only detected a few hundred lncRNAs. This observation is in line with the fact that the extracellular mRNA concentration in this sample type is low (Hulstaert *et al.*, 2020). Additionally, the blood plasma samples were not sequenced at high depth (3 million paired-end reads before duplicate removal), suggesting that results may improve when generating more reads.

For 10 to 25% of lncRNAs that were uniquely detected with total RNA-seq, no probes were present in the custom capture probe set because of a failure to satisfy probe design requirements. About half of the lncRNAs with at least one custom probe were still detected in some of the capture libraries but failed to reach the threshold in other replicates (and where therefore labeled as undetected in these libraries). While data was downsampled to the same read depth, increasing sequencing depth may solve this discrepancy. Some of the lncRNAs did not have probes complementary to the transcript regions that were detected with total RNA-seq. Incorporating additional probes against those regions could further improve performance, although this would require loosening probe design criteria, which may result in more non-specific hybridization and off-target capture. For the remaining lncRNAs, further optimization of the probe designs may be required to enable proper capture. Note that the custom capture library preparation is considerably more expensive than total RNA-sequencing. The price difference is mainly driven by the large custom probe set. Of note, probe cost could be substantially reduced when offered off-the-shelf or by transitioning from a discovery phase to a validation phase, including only those probes that target lncRNAs of interest. In this study, the stranded TruSeq RNA Exome Library Prep Kit was used for the custom capture approach, but other library prep methods would work too.

Taken together, we demonstrated that lncRNA capture sequencing is able to detect a more diverse repertoire of lncRNAs compared to standard total RNA sequencing, and increases coverage as well as reproducibility in both high-quality high input as well as fragmented and/or low input RNA samples.

## Acknowledgements

This work was supported by the Fund for Scientific Research Flanders (1226821N and 1S07416N to C.E.; 1133120N to E.H.; 11C1621N to A.M.) and the Special Research Fund (BOF) of Ghent University (BOF.D0C.2019.0047.01). This research is partly funded by “RNA-MAGIC” and “LNCCA” Concerted Research Actions of Ghent University (BOF19/G0A/008 and BOF16/GOA/023), by “Kom Op Tegen Kanker” (Stand Up To Cancer, the Flemish cancer society) and by the Foundation Against Cancer.

The authors wish to thank Kathleen Schoofs for preparing platelet-depleted plasma from blood samples and Kimberly Verniers for providing purified RNA of FFPE samples as well as her input on the custom lncRNA capture protocol.

We are very grateful to Gary Schroth, Scott Kuersten and Stephen Gross (Illumina) for providing a prototype set of lncRNA probes to evaluate the custom hybrid capture procedure.

## Supplemental Figures

**SFig 1:**
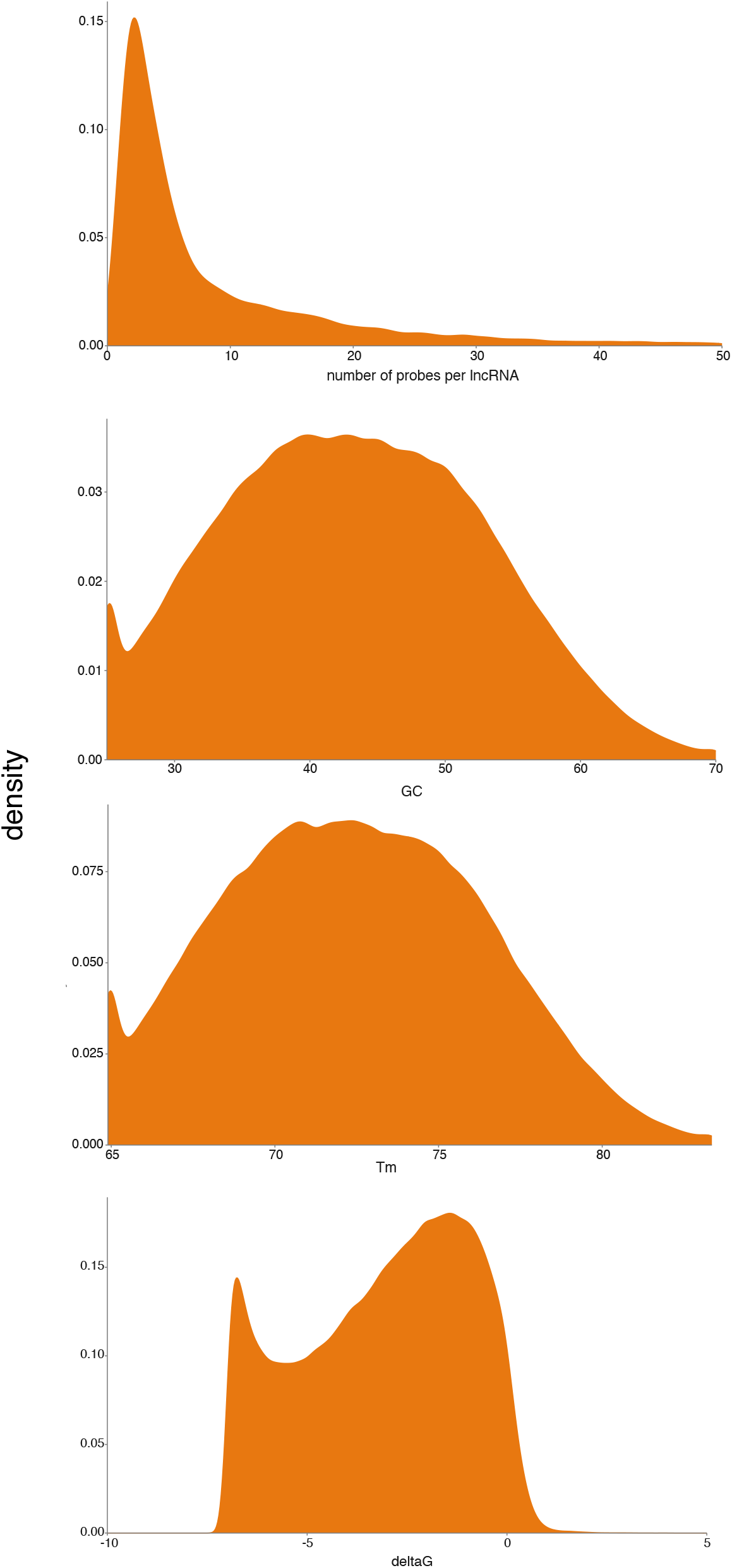
Distributions of number of probes per gene, GC%, melting temperature and ΔG of the selected probes.

**SFig 2:**
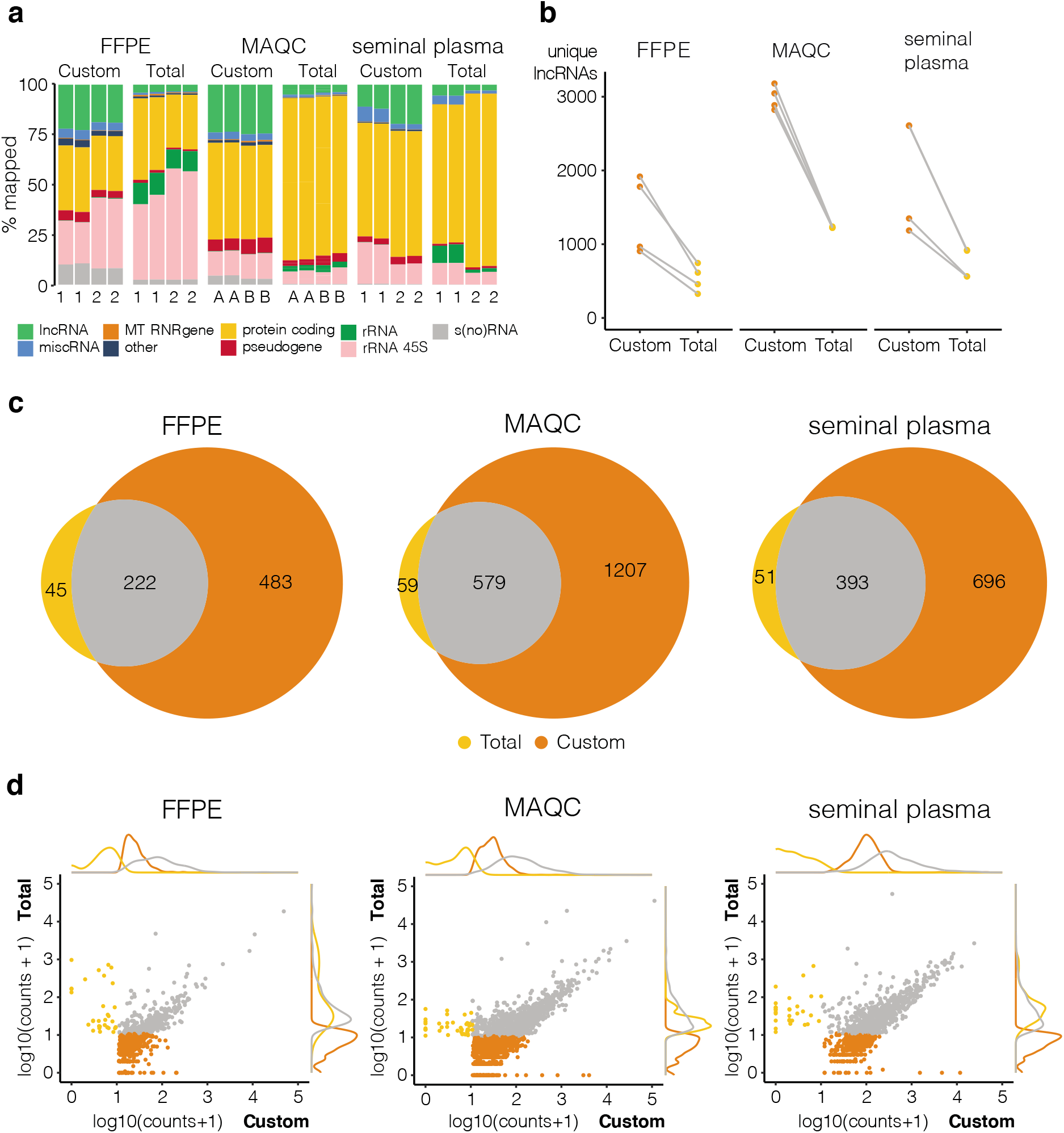
Custom capture sequencing (Custom) is able to detect more lncRNAs than total RNA-sequencing (Total) Quantification based on Ensembl v91 reference. a: RNA biotype distribution plot of mapped reads where 1 and 2 indicate the two different donors and A and B refer to MAQCA and MAQCB, respectively (lncRNAs: Ensembl lncRNAs; miscRNA: miscellaneous RNA, non-coding RNA that cannot be classified; MT RNR gene: mitochondrially encoded ribosomal RNAs; protein coding: protein coding RNA transcripts; pseudogene; rRNA (45S): (45S) ribosomal RNA; s(no)RNA: small nuclear/nucleolar RNA; ucgenes: unannotated cancer genes; other: T cell receptor genes, Immunoglobulin genes, TEC (To be Experimentally Confirmed) - regions with EST clusters that have polyA features that could indicate the presence of protein coding genes, vaultRNA - short non coding RNA genes that form part of the vault ribonucleoprotein complex; microRNAs; ribozymes); b: number of unique lncRNAs with at least 10 counts (filter threshold), data points from same donor are linked (grey lines); c: overlap between lncRNAs that are detected above threshold in all replicates of a certain library prep method, plots made with eulerr package (v6.1.0) in R; d: correlation and density plots of overlapping (grey) and specific lncRNAs for custom capture (orange) and total RNA-sequencing (yellow); lncRNAs below count threshold in both methods were left out.

**SFig 3:**
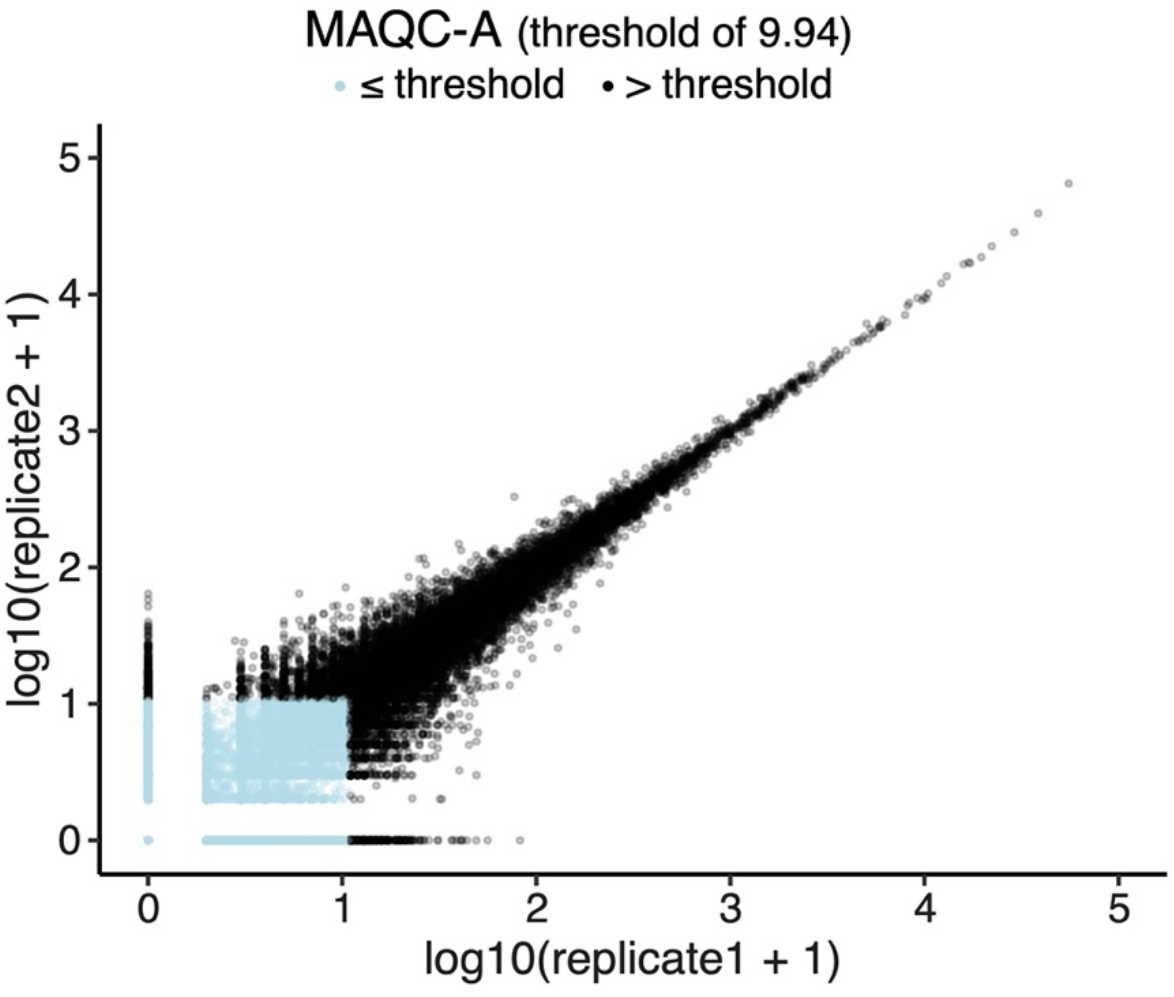
Reproducibility threshold is determined based on elimination of at least 95% of single positives between technical (library preparation) replicates. Single positives are detected (at least 1 count) in one replicate and not detected in the other replicate. Here, example of replicate correlation of MAQCA with total RNA sequencing is shown. Count threshold that filters out 95% of single positives is 9.94 (kallisto quantification leads to decimal counts). Data points that will be filtered with this threshold are in blue. Slope of linear model is 0.969, pearson correlation is 0.999.

**SFig 4:**
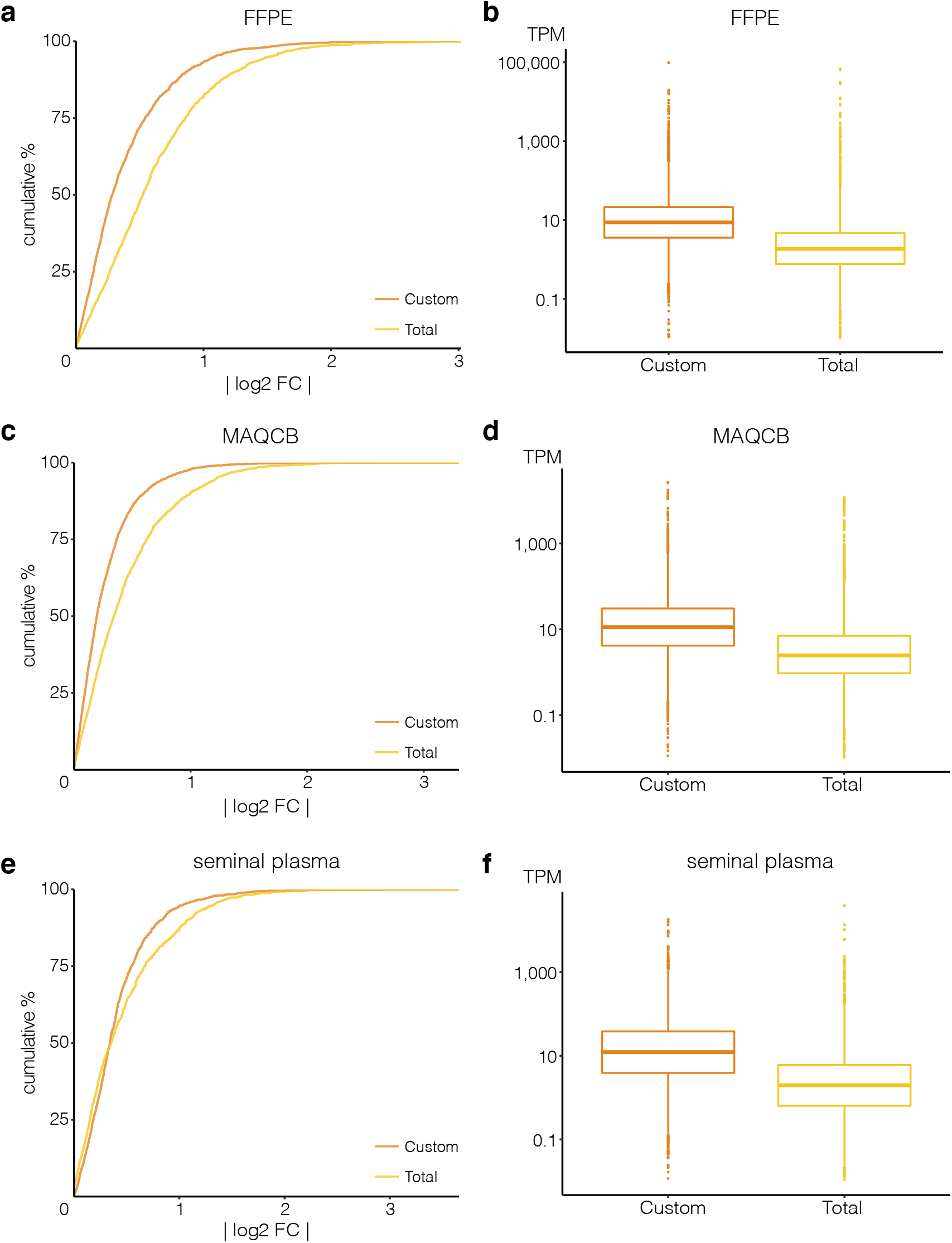
Custom capture seq (Custom) has a higher lncRNA count reproducibility and coverage than SMARTer Stranded Total RNA seq (Total). Cumulative distribution of absolute log2 fold changes between lncRNA counts in the two technical replicates are shown for (a) FFPE from donor 2, (c) MAQCB, and (e) seminal plasma from donor 1. Kolmogorov–Smirnov tests each time showed significant difference in distributions between Total and Custom (p-value < 0.001). Boxplot of corresponding transcripts per million (TPM) values of these lncRNAs are shown in (b) for FFPE, (d) for MAQCB, and (f) for seminal plasma.

**SFig 5:**
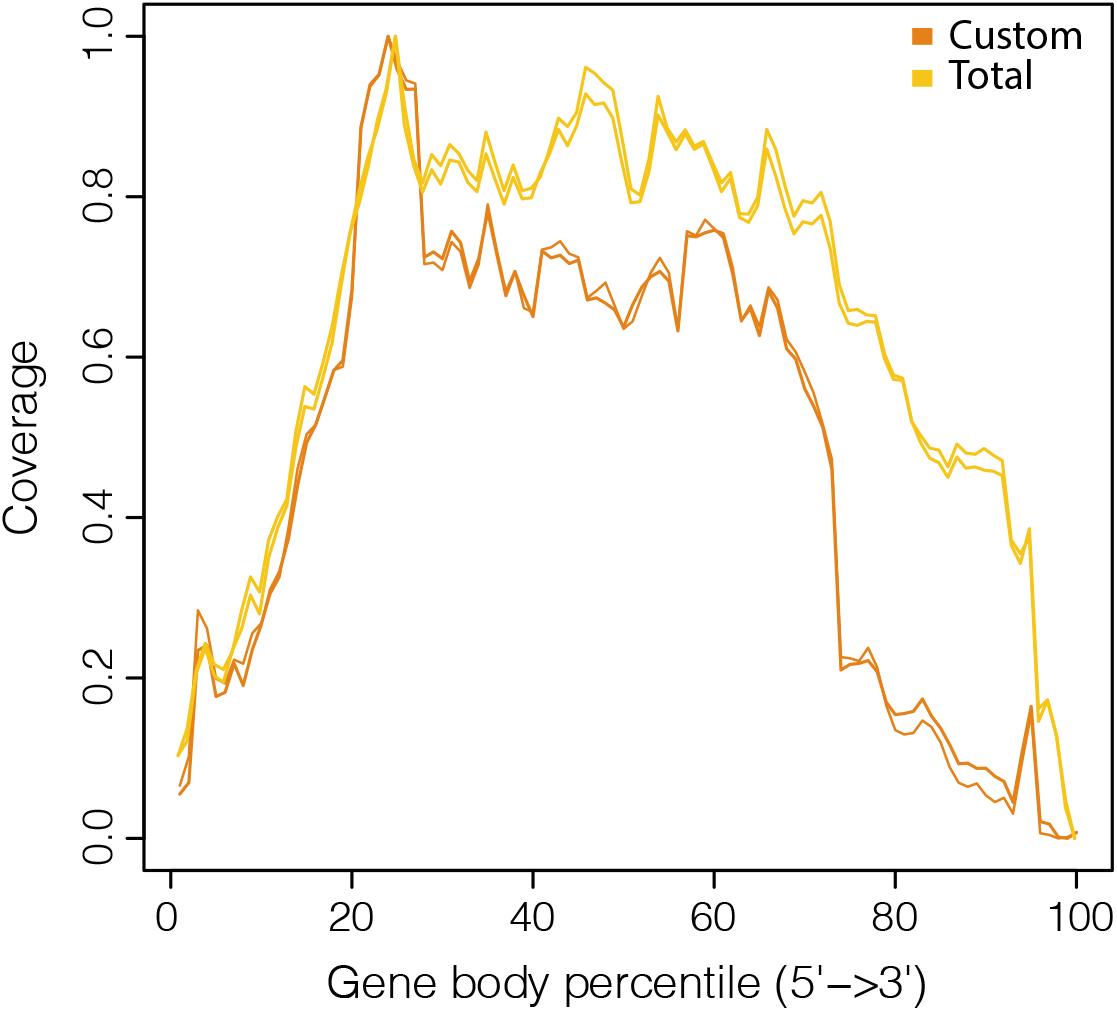
Distribution of gene body coverage shows quite stable coverage for majority of gene (lncRNA) body but drops towards the 5’ and 3’ ends. Distribution shown for both technical replicates of MAQCA. Custom: custom capture sequencing; Total: SMARTer Stranded Total RNA sequencing.

**SFig 6:**
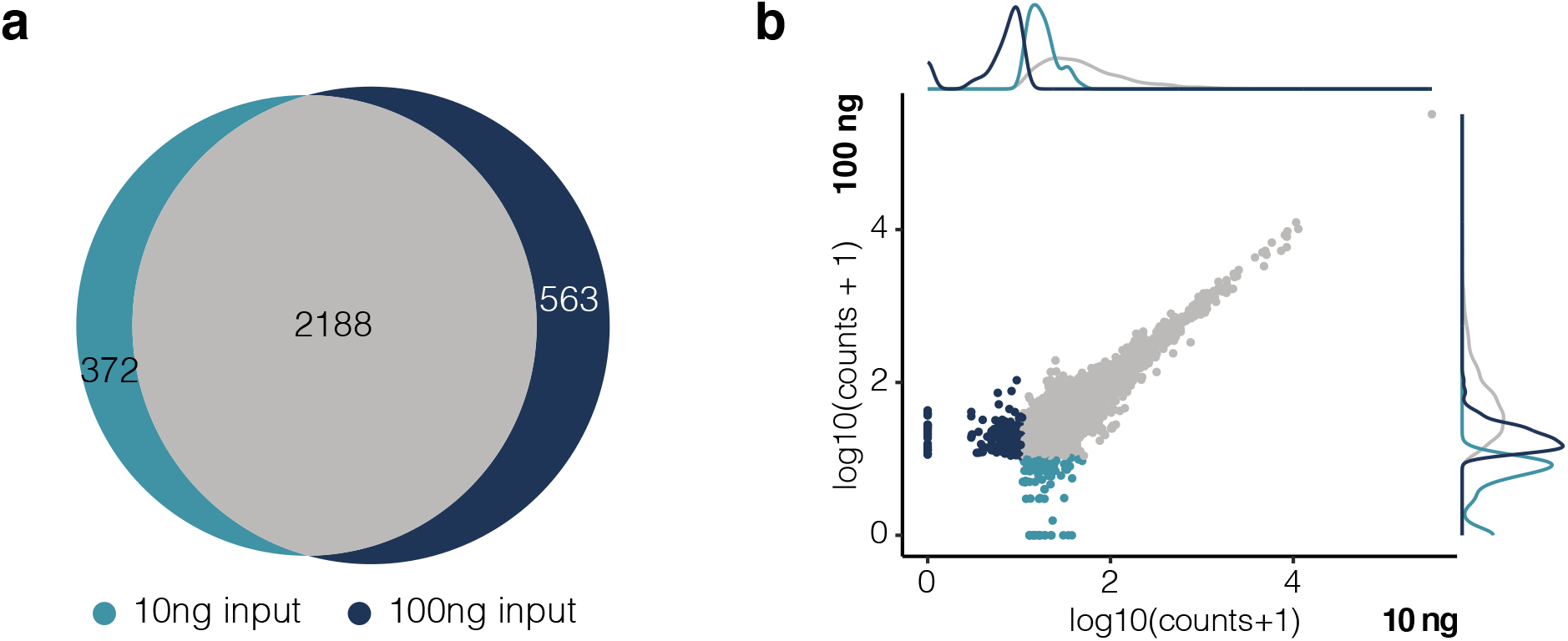
Impact of 10 vs 100 ng RNA input on SMARTer Stranded Total RNA sequencing results is limited. a: overlap between lncRNAs detected (> 10 counts) in MAQC with 10 vs 100 ng RNA input for library preparation; b: corresponding count correlation and density plot (counts shown for one technical replicate of MAQCA).

